# Kaleidoscope: A New Bioinformatics Pipeline Web Application for In Silico Hypothesis Exploration of Omics Signatures

**DOI:** 10.1101/2020.05.01.070805

**Authors:** Khaled Alganem, Rammohan Shukla, Hunter Eby, Mackenzie Abel, Xiaolu Zhang, William Brett McIntyre, Jiwon Lee, Christy Au-Yeung, Roshanak Asgariroozbehani, Roshni Panda, Sinead M O’Donovan, Adam Funk, Margaret Hahn, Jarek Meller, Robert McCullumsmith

## Abstract

**Background:** *In silico* data exploration is a key first step of exploring a research question. There are many publicly available databases and tools that offer appealing features to help with such a task. However, many applications lack exposure or are constrained with unfriendly or outdated user interfaces. Thus, it follows that there are many resources that are relevant to investigation of medical disorders that are underutilized.

**Results:** We developed an R Shiny web application, called Kaleidoscope, to address this challenge. The application offers access to several omics databases and tools to let users explore research questions *in silico*. The application is designed to be user-friendly with a unified user interface, while also scalable by offering the option of uploading user-defined datasets. We demonstrate the application features with a starting query of a single gene (Disrupted in schizophrenia 1, DISC1) to assess its protein-protein interactions network. We then explore expression levels of the gene network across tissues and cell types in the brain, as well as across 34 schizophrenia versus control differential gene expression datasets.

**Conclusion:** Kaleidoscope provides easy access to several databases and tools under a unified user interface to explore research questions *in silico*. The web application is open-source and freely available at https://kalganem.shinyapps.io/Kaleidoscope/. This application streamlines the process of *in silico* data exploration for users and expands the efficient use of these tools to stakeholders without specific bioinformatics expertise.

## Background

Increasing numbers of large biological datasets are being deposited into publicly available repositories (1). In conjunction, an ever-increasing number of bioinformatic tools are being developed to process, analyze and view a wide spectrum of biological datasets (2). However, the rapidly growing availability of databases and bioinformatic tools can be an impediment for scientists, often hampering discovery of these tools, especially for users who are not well versed in the bioinformatics field (3, 4).

We have observed that a large number of bioinformatics tools and databases that are relevant to psychiatric disorders are still relatively untapped. Even though some of these tools are well known, some researchers avoid utilizing them mainly due the sheer scale or the complexity of the user interface which can be overwhelming to novice users.

We addressed this issue by developing an interactive R Shiny web application, called Kaleidoscope. Kaleidoscope provides a platform for easy access to these resources via a user-friendly interface. Integrating multiple databases and tools in a single platform facilitates the users interactive exploration of research questions *in silico*. This approach to data exploration can lead to exciting observations that supplement existing hypotheses, generate new ones and possibly direct future studies (**Fig 1**) (5-8). This interactive exploratory data analysis platform is particularly targeted to a broader range of investigators who are not familiar with these tools. The platform solves the issue of outdated or complex user interfaces that impede many bioinformatics tools by presenting a simple and standardized user interface across the whole platform. Our web application utilizes application programing interfaces (APIs) to access databases to extract, harmonize, and present expression data using meaningful visualizations. An easy and fast process of examining datasets is vital to the concept of data exploration. The platform is also designed to accommodate user-provided datasets to better suit the user’s research interests. This in particular is a huge asset for investigators, as the process of finding, curating, formatting, and analyzing datasets is time consuming and requires trained individuals. Being aware of these challenges, we designed our application to minimize the process of data acquisition and formatting. This intended design allows researchers to primarily focus on *in silico* hypothesis exploration without spending unneeded time on the preceding steps. Moreover, as our user group grows and users upload their own curated datasets, we expect a sizable expansion of the number of datasets hosted in our platform. This promising feature will help grow the curated datasets found in the application in a user defined manner.

**Figure 1.**
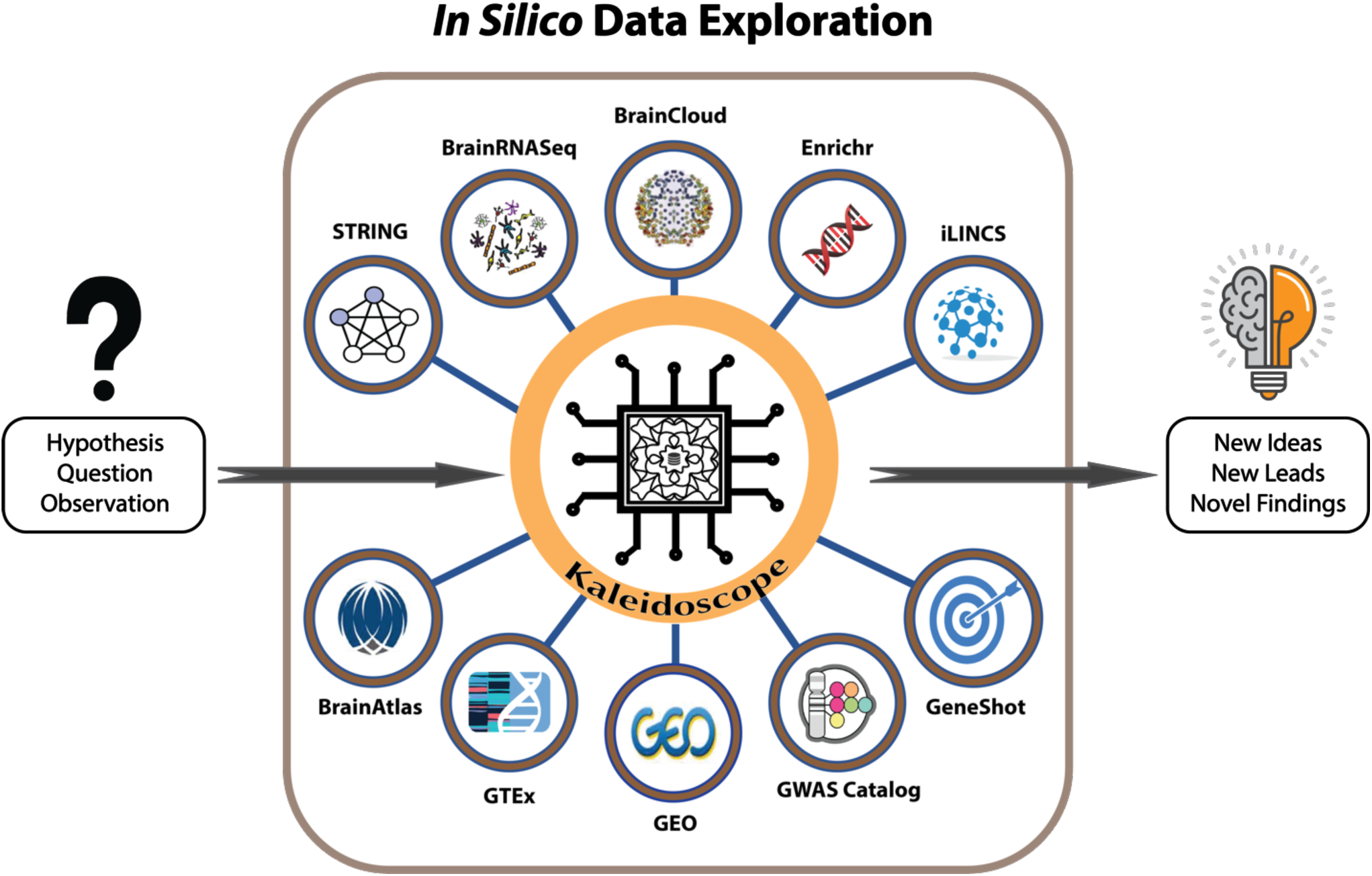
A workflow model for using Kaleidoscope to perform *in silico* exploratory data analyses. Starting with hypotheses and questions, Kaleidoscope integrates multiple platforms and tools to streamline the process of *in silico* data exploration. STRING (Search Tool for the Retrieval of Interacting Genes) provides protein-protein interaction networks. The Brain RNASeq database is used for cell specific gene expression levels in human and mouse brain tissues. BrainCloud is used to extract gene expression patterns in the brain across the lifespan of healthy humans. Enrichr is integrated to perform gene set enrichment analyses. iLINCS (Integrative Library of Integrated Network-Based Cellular Signatures) is queried to search for gene knockdown transcriptional signatures and generate L1000 signatures that can be further explored using the iLINCS platform. BrainAtlas is used to assess gene expression levels in different brain cell types. GTEx (Genotype-Tissue Expression) is used to explore tissue specific gene expression patterns. GEO (Gene Expression Omnibus) is utilized to extract and curate previously published differential gene expression datasets. GWAS Catalog (Genome-Wide Association Study Catalog) is searched for previously published genome-wide association studies. The GeneShot tool is used to text mine published studies to associate search terms and genes.

## Implementation

Kaleidoscope is an R Shiny web application that integrates multiple databases that are relevant to psychiatric disorders; Brain RNA-Seq, BrainCloud, BrainAtlas, BrainSpan, STRING, iLINCS, Enrichr, GeneShot, and GWAS Catalog, as well as over 200 (and counting) disease-related differential gene expression datasets.

### Brain RNA-Seq

The Brain RNA-Seq provides insight on gene expression of isolated and purified cells from human and mouse cortical tissue (9, 10). Our application utilizes the Brain RNA-Seq database to incorporate a searchable database of cell-specific mRNA expression in rodent and human brain tissues. The Brain RNA-Seq database has its own web interface, but it’s limited to only searching one gene at a time. Our platform permits inquiry of multiple genes simultaneously, while also displaying figures that represent the ratio of expression across the different cell types that are present in the database. This module in our platform can serve as a first step for assessing expression levels of target genes in different brain cells to understand their cell-specific function and role, while also highlighting the differences of profiles between the two species.

### Search Tool for the Retrieval of Interacting Genes (STRING)

STRING is a popular database of known and predicted protein-protein interaction (PPI) networks (11). These interactions represent either physical or functional associations between proteins. This module in our application can be used to grow a gene of interest into a network by querying for PPI networks. Through a connection to the STRING API, the user has the option to specify the stringency of the predicated associations (by choosing the appropriate score cutoff), choosing the desired number of connected proteins, and the desired organism. Our application efficiently communicates the complex results from STRING by displaying the PPI network together with a figure legend, providing context for the different edges in the network. Moreover, a table is displayed that lists all of the proteins in the network with a brief description to each corresponding protein, as well as the values of each scoring method that STRING uses to calculate the combined scores (12).

### BrainCloud

The BrainCloud database was developed through a collaboration between the Lieber Institute and The National Institute of Mental Health (NIMH) to give a global insight of the role of the human transcriptome in cortical development and aging (13). Using the data generated by BrainCloud, we can observe the patterns of expression in the brain of our target genes across the lifespan of healthy humans. This data is used to explore the genetic control of transcription of our targets during development and aging using samples from the prefrontal cortex.

### Allen Brain Map

We utilized the cell type database of the Allen Brain Map which examines the transcriptional profile of thousands of single cells with RNA-Seq (14). Currently our platform has access to RNA-Seq data, generated from intact nuclei derived from frozen human brain specimens, to survey cell type diversity in the human middle temporal gyrus (MTG). In total, 15,928 nuclei from 8 human tissue donors ranging in age from 24- 66 years were analyzed (14). Analysis of these transcriptional profiles reveals approximately 75 transcriptionally distinct cell types, subdivided into 45 inhibitory neuron types, 24 excitatory neuron types, and 6 non-neuronal types (14). A list of target genes can be queried across all of these clusters of cell subtypes to examine their expression levels or enrichments. A heatmap with unsupervised hierarchical clustering is displayed to visualize the differences of gene expression across the target genes and also across the different cell subtype clusters. Additionally, the user may get results about whether genes are enriched in one cell subtype cluster versus another by comparing the proportions of expression across all cell subtypes, with an adjustable difference cutoff.

In addition, the BrainSpan atlas is integrated into the platform which complements the BrainCloud database by exploring the transcriptional mechanisms involved in human brain development, but with specific transcriptional profiling of different brain regions (15). Currently, 10 brain regions are included in Kaleidoscope. Data are displayed as Reads Per Kilobase Million (RPKM) expression values.

## GTEx

The Genotype-Tissue Expression (GTEx) database contains data of tissue-specific gene expression and regulation. The database is constructed based on samples that were collected from 54 non-diseased tissues from almost 1000 individuals, primarily for molecular assays including whole genome and transcriptome sequencing (16). Our application allows the user to input a list of genes to query their expression levels across all of the tissues that were analyzed in the GTEx database. The results display the median TPM (Transcripts Per Kilobase Million) of each gene in the list presented as heatmap, with options of unsupervised hierarchical clustering or log transformation. This module is very helpful in capturing the tissue-specific regulation of expression of the user’s genes of interest.

Moreover, the GTEx database contains eQTL (expression quantitative trait loci) mapping for almost all studied tissues to identify variant-gene expression associations. Our platform has access to eQTL mapping of tissues relative to our field, including samples from 5 different brain regions.

### Lookup Replication Studies

Previously published, peer-reviewed, and publicly available transcriptomics and proteomics datasets were carefully curated and selected to probe for patterns of differential gene expression between healthy subjects and diseased/perturbed samples. These datasets were analyzed using well-established differential expression analysis R packages. Specifically, we used Limma for microarray datasets, and EdgeR/DESeq2 for RNASeq datasets (17-19).

The curated datasets cover several substrates, including stem cells, postmortem brain, and animal models. At present the application has over 200 datasets grouped under different modules. The modules currently available are schizophrenia, depression, antipsychotics, dopamine signaling, insulin signaling, bipolar disorder, Alzheimer’s disease, aging, microcystin, and coronavirus. These modules are also being expanded as we integrate additional datasets. In addition, users have the ability to upload their own curated datasets into the application, thus expanding the different diseases hosted in the platform. Loaded datasets are automatically harmonized by calculating the empirical cumulative probabilities of the log2 fold change values of genes within each dataset (20).

Kaleidoscope displays the results for this section of the software across multiple tabs. The “Results” tab shows a table where each row represent a gene from the input list of genes and each column represent a dataset from the selected datasets. The values in the table are the log2 fold-change values and p-values. The “Lookup Graph” tab shows a figure to visualize the results from the table from the “Results” tab with additional information to help the user quickly navigate the results. The x-axis of the figure represents genes and the y-axis represents proportion values of number of “hits,” log2 fold change values that passed a cutoff threshold which can be adjusted by the user. These values represent the number of hits over the number of datasets for each target gene, excluding datasets that have missing values. The colors represent the number of datasets which are found for each gene. The “Heatmap” tab displays an interactive heatmap representing log2 fold change values (Log2FC), fold change values (FC), and standardized scores based on the calculations of empirical cumulative probabilities (ECDF). These heatmaps provide unsupervised hierarchical clustering for both genes and datasets. The clustering allows patterns of similar changes of expression across the list of genes and the datasets that were selected to be identified. The “Correlation” tab in this section shows a visualization of the concordance scores matrix calculated using either Pearson or Spearman correlation analysis based on the log2 fold change values. The user has the option to calculate the concordance scores based on the input list of genes or the full list of genes in the datasets. For each correlation analysis between two datasets, the test is applied using only the genes that were found in both datasets. The colors on the figure denotes the direction of the concordance scores, where red represents negative correlation (high discordance) and blue represents positive correlation (high concordance). The “References” tab shows a table with brief descriptions of each dataset and their references.

Alternatively, instead of inputting a list of genes, the user has the option to query the commonly differentially expressed genes across multiple datasets by sorting the list of genes based on their absolute log2 fold change values within each dataset and extracting the top list of genes. An additional tab is shown when the user selects this type of analysis; it displays a heatmap representing the overlapping hits across the queried datasets. The user can adjust the parameters of this analysis by selecting the desired number of extracted genes from the datasets and the final number of genes that have the most overlap across the selected datasets.

This section of the application helps the user to “lookup” expression patterns from publicly available and curated datasets to find any interesting patterns of gene expression changes. Seeing “hits” or above average gene expression differences across multiple datasets is a strong indication that a gene or a panel of genes is involved in the disease process. Additionally, it can be used to complement a user’s own transcriptome study that is related to any of the existing diseased/perturbed modules to conduct quick and easy lookup, replication, or confirmation studies (5).

### The Library of Integrated Network-Based Cellular Signatures (LINCS)

The LINCS database is a large multi-omics profiling database. For transcriptional datasets, it utilizes the L1000. The L1000 is a gene-expression profiling assay based on the direct measurement of a reduced representation of the transcriptome (978 “landmark” genes) under different perturbations; gene knockdown, gene overexpression, and drug treatments (21). Kaleidoscope provides the option to generate L1000 signatures by extracting the 978 genes and averaging the log2 fold change values across the selected datasets. The user is then able to download the L1000 signature as a tab delimited file, and upload it to the integrative LINCS (iLINCS) portal to perform perturbagen connectivity analysis. iLINCS is a web platform developed to explore and analyze LINCS signatures and is mainly used to perform *in-silico* drug discovery analysis (22).

In our application, the user can also query gene knockdown signatures by inputting a list of genes and will be connected to iLINCS API. The results are displayed as a table with the total number of knockdown signatures found per gene. Also, another table is displayed with more information on these gene knockdown signatures by specifying the gene knockdown signatures ID, cell line, and a direct link to the signature web page on the iLINCS web portal.

### Genome-Wide Association Study (GWAS) Catalog

The GWAS Catalog is supported by the National Human Genome Research Institute (NHGRI). The GWAS Catalog database is the largest curated collection of all published genome-wide association studies (23). In our application, the user can query the GWAS Catalog database by inputting a gene or a list of genes. The results are displayed as a table with all significant single-nucleotide polymorphisms (SNPs) mapped to the input genes. The table will show the gene name, SNP ID (rs ID), chromosome position, studied disease or phenotype, and a direct link to the study. In addition, the type of SNP will be shown as intergenic variant, 3 Prime UTR variant, regulatory region variant, etc. Two additional tabs display distinct figures: the first displaying the top overlapping disease/phenotype traits, the second an interactive sankey graph to represent the flow rate between the list of genes, SNP type, disease/phenotype trait.

### Enrichr and GeneShot

Enrichr and GeneShot are two tools that were developed by The Ma’ayan Laboratory, Icahn School of Medicine at Mount Sinai (24, 25). Enrichr is a tool for performing gene set enrichment analysis across many databases (24). GeneShot is a search engine for search terms and genes mentions based on arbitrary text queries of published literature (25). Kaleidoscope utilizes these tools’ APIs to integrate them across Kaleidoscope for easy access to gene set enrichment analyses.

## Discussion

To demonstrate the application’s features, we ran an example session starting with a single gene of interest. The purpose of this demonstration is to highlight some of the rich information and data that can be extracted using Kaleidoscope. We picked the Disrupted in schizophrenia 1 (DISC1) gene as the starting gene of interest. DISC1 has emerged as a strong candidate gene underlying the risk for major mental disorders (26, 27). We expanded our single gene into a network using the STRING tab, by selecting human as the desired species and using 500 as the score cutoff as well as limiting the desired number of nodes to 25 (**Fig 2A**). A table is generated that displays a brief description of each gene, and association scores under the different criteria (experimental database, co-expression, text mining etc.) (**Supplementary Table 1**). Using this list of genes from the PPI network, we data mined the other available databases in Kaleidoscope. We used the GeneShot tab to search the relevance of schizophrenia and our PPI network across previous publications. In **Fig 2B** the total publications that mention each gene and the proportions of these publications that also mention schizophrenia is shown. As expected, DISC1 has a high proportion of publications that were linked to schizophrenia. Interestingly, some genes in the network appeared to be understudied with regards to their association to schizophrenia (Serine racemase (SSR), Oligodendrocyte transcription factor 2 (OLIG2), Glycogen synthase kinase-3 beta (GSK3B)).

**Figure 2.**
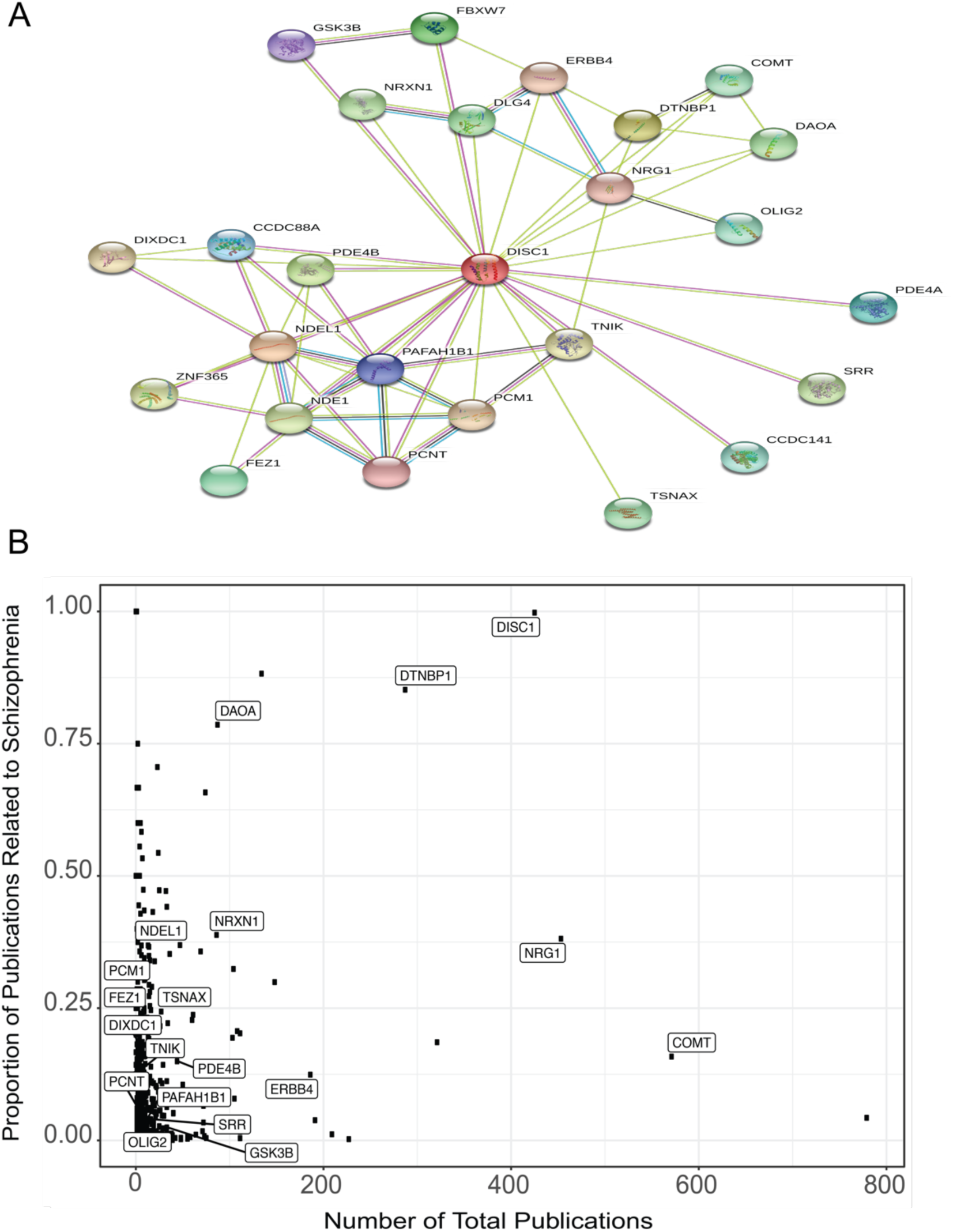
Protein-protein interaction (PPI) network for the DISC1 protein and mentions in the schizophrenia literature. A) The PPI network generated from the STRING (Search Tool for the Retrieval of Interacting Genes) database for DISC1 and its 25 close interactors (0.500 was used as the minimum required interaction score). B) A scatterplot to represent the associations of the DISC1 gene set from the PPI network with schizophrenia (number of papers that mention both the gene name and schizophrenia over the number of total number of papers that mention that gene).

Next, the Brain RNA-Seq tab was used to explore DISC1 cell type specific gene expression. Fragments Per Kilobase of transcript per Million mapped reads (FPKM) for DISC1 in both human and mouse are shown for multiple cell types (neurons, astrocytes, endothelial cells, oligodendrocyte progenitor cells, etc.) (**Fig S1**). Additionally, we queried the Brain RNA-Seq tab to look at cell-type specific gene expression, using the whole network and selecting the option of multiple targets input. The gene expression figures for each gene are displayed. Boxplot and proportion figures are also displayed when using the multiple target option. We observed higher levels of expression for this network in neurons and oligodendrocytes (**Fig 3A-B**). Oligodendrocytes have been linked to schizophrenia in regards to myelin dysfunction and neurocircuitry abnormalities (28).

**Figure 3.**
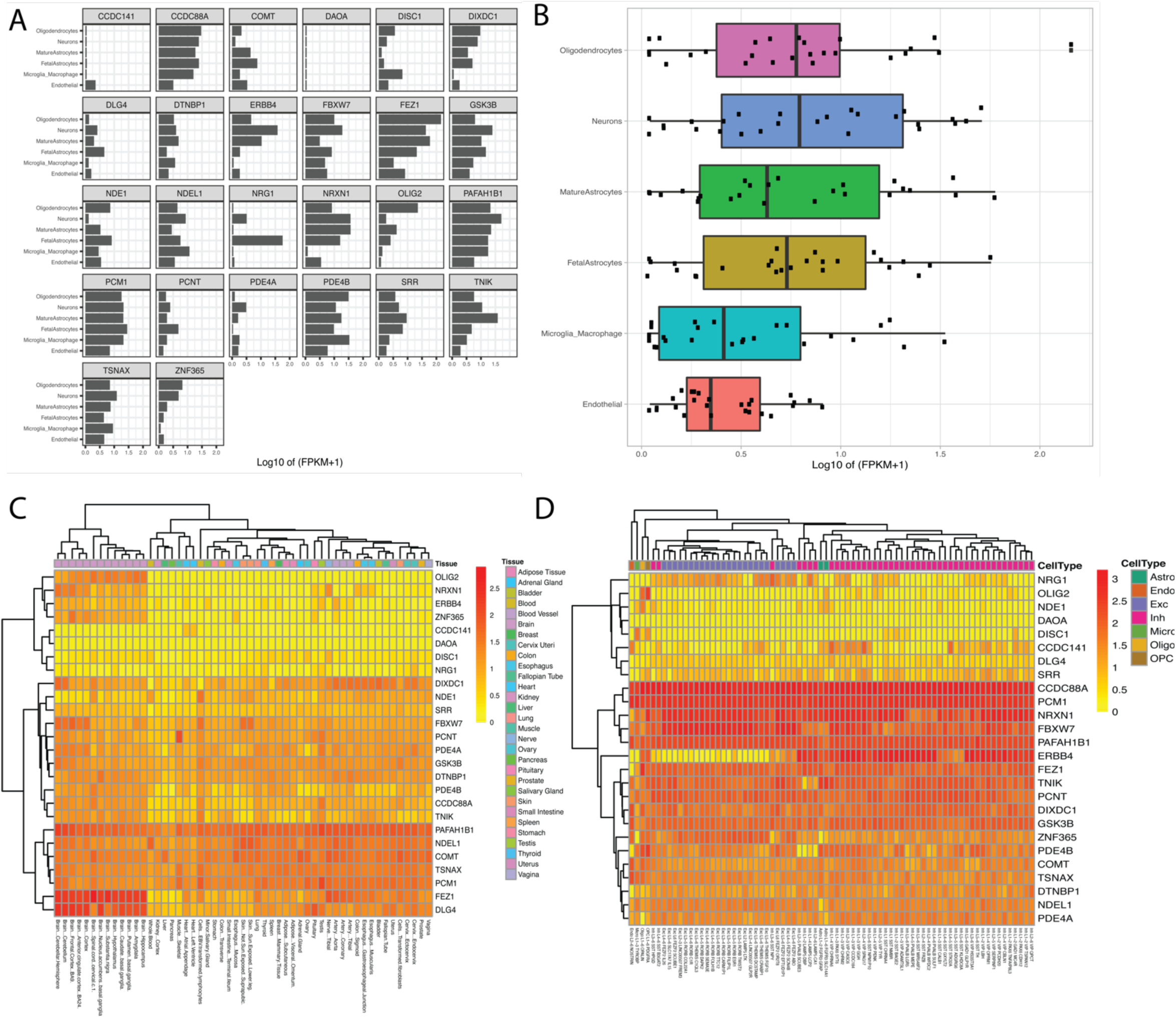
Plots and heatmaps to represent transcription enrichments across different cell types and tissues of the DISC1 protein interactors. A) Results from the Brain RNA-Seq. Each facet represents a gene, and the values show expression levels (log2(FPKM+1)) across the 5 different cell types in human brain tissue. B) Boxplots to show the distribution of gene expression levels for the full list of genes. C) A heatmap to highlight the difference of gene expression levels in different tissues from the GTEx database. D) Heatmap to show the gene expression levels in the different cell types from the BrainAtlas with unsupervised hierarchical clustering

We continue to explore the expression levels of our target genes in different cell types, this time using the BrainAtlas tab. Inputting our list of genes in that tab yields a table and a gene expression heatmap. The table displays the range of count per million (CPM) values for each gene across all of the different cell types, using all the sub- clusters within each cell type. A heatmap is also generated with unsupervised hierarchical clustering for both genes and cell type sub clusters (**Fig 3C**). GTEx was also queried to observe tissue specific gene expression levels. Samples from brain tissues clustered together, suggesting a similar pattern of expression of gene networks in the brain compared to other tissue types. Also, an enrichment of gene expression in brain tissues has been observed for many of the genes in our network, including OLIG2, NRXN1 and DLG4 (**Fig 3D**). We then used the lookup tab to search relative gene expression changes of the network of genes across different schizophrenia transcriptional datasets (29-42). Several figures and tables are displayed, most importantly a heatmap of log2 fold change values and table of gene expression changes across the different datasets (**Fig 4A, Supplementary Table 2**). The results indicate a clear dysregulation in expression for our network of genes, consistent with previous findings of their association with schizophrenia (43-45). Also, a correlation matrix figure is shown based on the list of the genes from the DISC1 network and paired based on the selected datasets (**Fig 4B**). Finally, the iLINCS tab was utilized to search for knockdown signatures of the list of genes from the network and generate a table of signature IDs, cell line, and direct link to the signature iLINCS webpage for further exploration (**Supplementary Table 3**).

**Figure 4.**
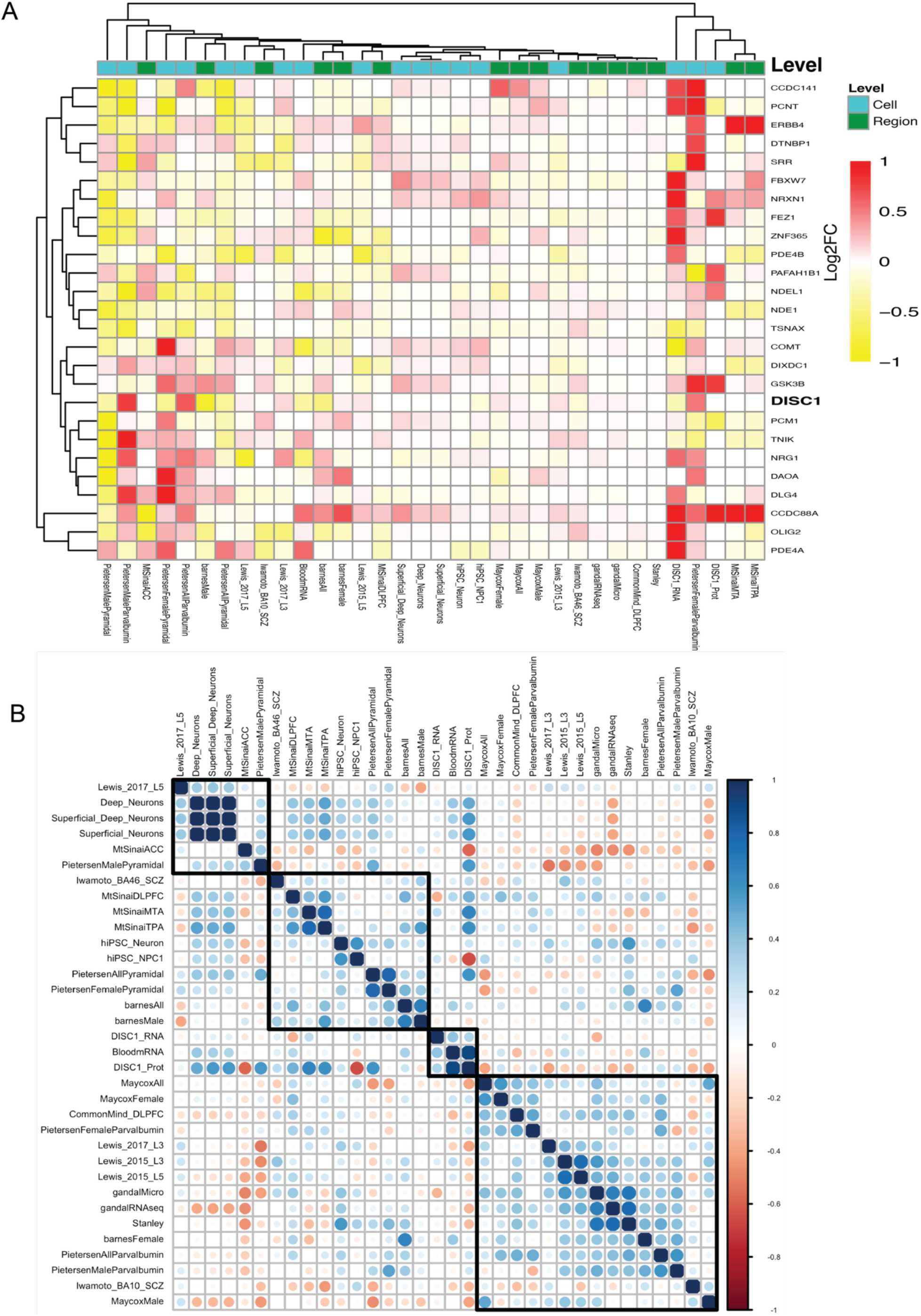
Differential gene expression of DISC1 protein interactors across 34 schizophrenia datasets. A) Heatmap with log2 fold change values between cases and controls across curated schizophrenia datasets. The datasets are grouped by their sample level (Region; samples taken from tissues, Cell; samples taken from pools of isolated cells). B) Correlation analysis of log2 fold change values of the DISC1 protein interactors between the schizophrenia datasets (Spearman correlation). Blue represents high concordance and red represents high discordance.

## Conclusion

As demonstrated here, Kaleidoscope provides an integrated platform to perform *in silico* data exploration of transcriptional signatures. The ability to efficiently explore DISC1 related signatures across all of these databases and tools offers insights on its protein-protein interactors and their regulation patterns across schizophrenia transcriptional datasets. Kaleidoscope has been recently utilized to perform *in silico* replication analyses of publicly available datasets, highlighting its utility (5-8). Our platform was used to supplement findings of bioenergetic gene expression dysregulation, and to explore adenosine system dysregulation in schizophrenia (5, 6). It was also used to explore abnormal regulators of protein prenylation in schizophrenia (7). Most recently, Kaleidoscope was utilized to investigate alteration of glutamate transporter interacting proteins in schizophrenia, major depression, and amyotrophic lateral sclerosis (8). Taken together, these examples show the advantage of having a platform that streamlines the process of *in silico* data exploration, making bioinformatics tools accessible to a wider range of users to test and investigate scientific questions and findings *in silico*.

## Availability and requirements

Project name: Kaleidoscope

Webpage: https://kalganem.shinyapps.io/Kaleidoscope/

Project home page: https://github.com/kalganem/kaleidoscope

Operating system(s): Platform independent

Programming language: R

Other requirements: e.g. Dependent R packages

License: GNU GPL.

Any restrictions to use by non-academics: none

## List of abbreviations

API: Application programing interface
PPI: Protein-protein interaction
STRING: Search Tool for the Retrieval of Interacting Genes
MTG: middle temporal gyrus
RPKM: Reads Per Kilobase Million
TPM: Transcripts Per Kilobase Million
eQTL: Expression quantitative trait loci
SNP: Single nucleotide polymorphism
LINCS: The Library of Integrated Network-Based Cellular Signatures
iLINCS: Integrative LINCS
GWAS: genome-wide association study
UTR: untranslated region
FPKM: Fragments Per Kilobase of transcript per Million mapped reads
CPM: count per million

## Declarations

### Ethics approval and consent to participate

Not applicable

### Consent for publication

Not applicable

### Availability of data and materials

The datasets generated and/or analyzed during the current study are available in the GitHub repository, https://github.com/kalganem/kaleidoscope/tree/master/data The application is available at https://kalganem.shinyapps.io/Kaleidoscope/

### Authors’ contributions

KA developed and designed the software. KA and RM wrote the manuscript. RS, SO, AF, MH, JM and RM provided feedback on the application. KA, HE, MA, XZ, WM, JL, CA, RA, and RP curated the datasets. All authors read and approved the final manuscript.

**Supplementary Table 1:**
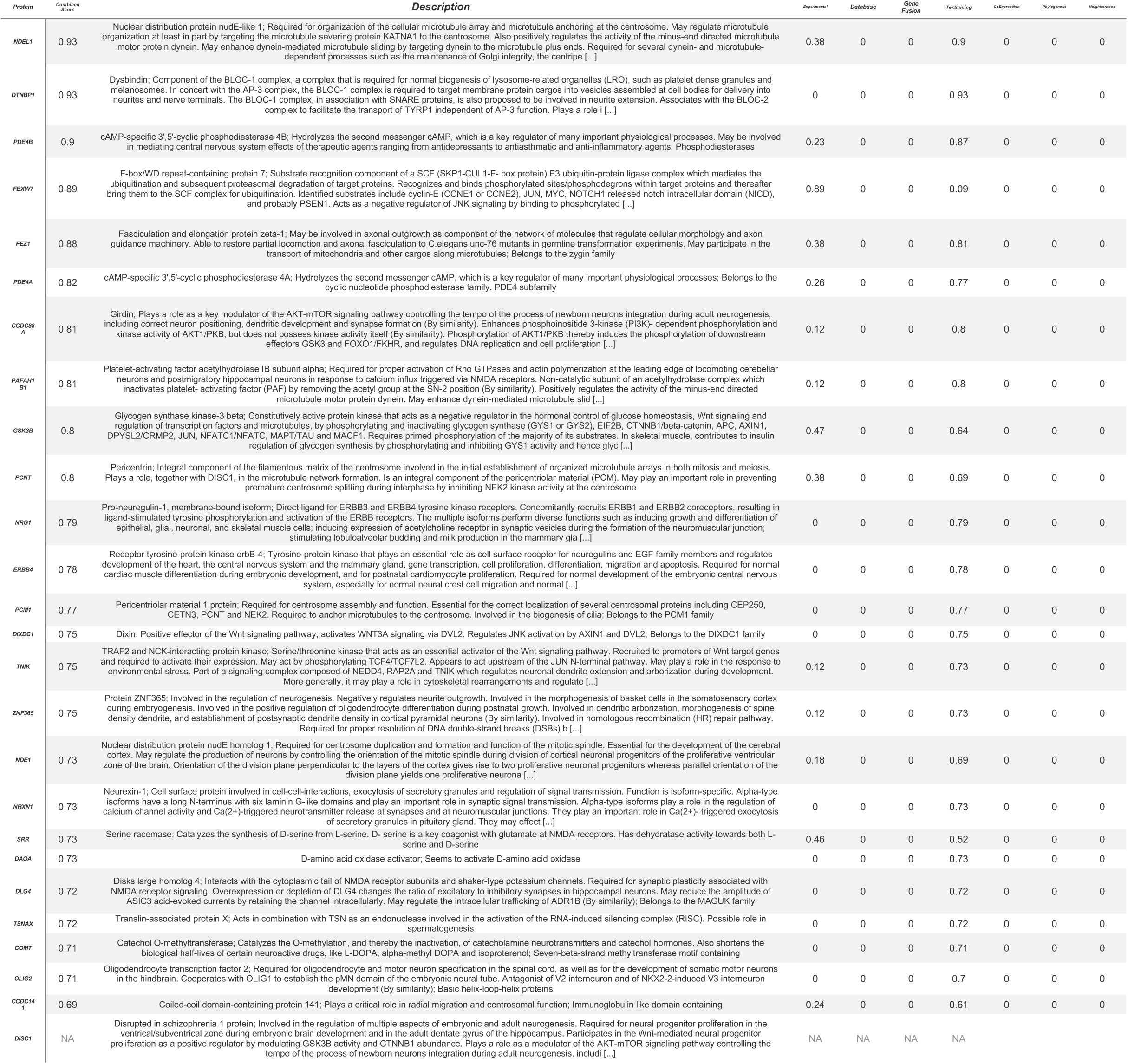
STRING protein-protein association network for the DISC1 protein. The table displays a brief description of each protein, and association scores under the different criteria. The combined score is computed based on the assumption of independence of the other scores (12).

**Supplementary Table 3:** iLINCS gene knockdown signatures table. A tabular summary displaying the number of gene knockdown signatures found in iLINCS for each gene in the protein-protein interaction (PPI) network for the DISC1 protein.

| GENE | KNOCKDOWN SIGNATURES |
| --- | --- |
| **GSK3B** | 15 |
| **COMT** | 11 |
| **ERBB4** | 11 |
| **FEZ1** | 11 |
| **PAFAH1B1** | 11 |
| **NRG1** | 10 |
| **PCM1** | 10 |
| **TNIK** | 10 |
| **ZNF365** | 8 |
| **DISC1** | 2 |
| **DIXDC1** | 2 |
| **NRXN1** | 2 |

## Acknowledgements

Not applicable

